# Synchro-PASEF allows precursor-specific fragment ion extraction and interference removal in data-independent acquisition

**DOI:** 10.1101/2022.11.01.514654

**Authors:** Patricia Skowronek, Florian Krohs, Markus Lubeck, Georg Wallmann, Ericka Itang, Polina Koval, Maria Wahle, Marvin Thielert, Florian Meier, Sander Willems, Oliver Raether, Matthias Mann

## Abstract

Data-independent acquisition (DIA) methods have become increasingly popular in mass spectrometry (MS)-based proteomics because they enable continuous acquisition of fragment spectra for all precursors simultaneously. However, these advantages come with the challenge of correctly reconstructing the precursor-fragment relationships in these highly convoluted spectra for reliable identification and quantification. Here we introduce a scan mode for the combination of trapped ion mobility spectrometry (TIMS) with parallel accumulation – serial fragmentation (PASEF) that seamlessly and continuously follows the natural shape of the ion cloud in ion mobility and peptide precursor mass dimensions. Termed synchro-PASEF, it increases the detected fragment ion current several-fold at sub-second cycle times. Consecutive quadrupole selection windows move synchronously through the mass and ion mobility range, defining precursor-quadrupole relationships. In this process, the quadrupole slices through the peptide precursors, which separates fragment ion signals of each precursor into adjacent synchro-PASEF scans. This precisely defines precursor – fragment relationships in ion mobility and mass dimensions and effectively deconvolutes the DIA fragment space. Importantly, the partitioned parts of the fragment ion transitions provide a further dimension of specificity via a lock and key mechanism. This is also advantageous for quantification, where signals from interfering precursors in the DIA selection window do not affect all partitions of the fragment ion, allowing to retain only the specific parts for quantification. Overall, we establish the defining features of synchro-PASEF and explore its potential for proteomic analyses.

## INTRODUCTION

Data independent acquisition (DIA) has shown enormous promise in recent years because of its signature advantage that every precursor is guaranteed to be fragmented once within every DIA acquisition cycle (1). Although this does not guarantee that every peptide is identified and quantified in every run, it goes a long way to make data sets more reproducible. Importantly, the fragment ions in DIA have elution profiles just like precursors, which dramatically improves to the quality of the fragment peaks. Frequently used software tools such as DIA-NN (2, 3) or Spectronaut (4) match peptides from a library into each of the DIA runs. These libraries were traditionally acquired experimentally with deep data dependent acquisition (DDA)-based measurements of the proteome of interest, but are now often generated directly from the DIA data (5) or in silico from the entire proteome using deep learning (6–11). Given these advantages very deep and quantitatively accurate data sets can now routinely be generated by DIA.

Despite these rapid developments, many researchers still prefer data dependent acquisition – often because only a single precursor is targeted for fragmentation, leading to simpler fragmentation spectra and more straightforward peptide identifications that may appear more trustworthy.

Following the original implementation of the SWATH acquisition scheme on the Sciex platform by Aebersold and colleagues (1, 12) and then on the Orbitrap instruments, where DIA first demonstrated in-depth proteome coverage (4), we have recently combined DIA with the Bruker timsTOF (13). In this instrument, peptides are confined in a trapped ion mobility spectrometry (TIMS) device by the opposing forces of a gas flow and an electric field and then released from the trap by lowering the electric field strength. Collisional cross sections and molecular weights of charged peptides are highly correlated, which is the basis of the ‘parallel accumulation − serial fragmentation’ (PASEF) principle, which boosts fragmentation rates in DDA more than ten-fold at no loss of sensitivity (14, 15). For the DIA mode, we very recently described a refined acquisition scheme that simultaneously optimizes the ion mobility and mass ranges for best precursor coverage (16). However, in these schemes, the quadrupole selection window remains fixed for each segment of the acquisition cycle.

We reasoned that a synchronous movement of the quadrupole selection window with the ion mobility scan would better utilize the trapped ion cloud. Furthermore, the rapidly advancing quadrupole window across the mass and ion mobility ranges could lead to much more defined precursor – fragment relationships. Although this scan mode is reminiscent of the one-dimensional movement of the quadrupole in techniques such as SONAR (17, 18) and Scanning SWATH (19), there are important conceptional differences. Simply scanning the m/z range in a timsTOF would imply very wide selection windows to cover the extent of the ion cloud in the mass dimension and would fail to capture most precursors if it was not synchronized with the release of the ions from the TIMS device.

We therefore set out to develop a synchronized PASEF scan mode with very high speed, so as to cover the entire precursor mass range of interest ideally in a single TIMS ramp. We first investigate the analytical properties of ‘synchro-PASEF’ on simple protein mixtures. Our results demonstrate that it makes efficient use of the ion population for fragmentation, even with very short acquisition cycles. We also find that peptide precursors are nearly universally ‘sliced’ in the ion mobility dimension by adjacent synchro-PASEF scans and describe how this enables the extraction of pure fragmentation spectra and mitigates interference for quantification.

## EXPERIMENTAL PROCEDURES

### Development of an extremely fast and synchronized scan capability

Previously in dda- and dia-PASEF, the analytical quadrupole was operated statically, i.e. stepped through a defined number of isolation windows per TIMS ramp. While the quadrupole changed its position, the signal was not recorded, limiting the number of quadrupole steps. In synchro-PASEF, the quadrupole selection window changes incrementally and is constantly synchronized to the TIMS elution. To acquire the complete signal, we routed the digital analog converters (DACs) of the quadrupole RF-amplitude, bias-DC, and resolution-DC to the same field programmable gate array (FPGA) of the instrument controller that controls the TIMS device. We also implemented the mass-to-set-value calculations based on the quadrupole calibration in the FPGA. Translation of the setpoint isolation masses and widths to the DAC values is implemented on the firmware level.

Together, this results in a more accurate correlation of the quadrupole with the TIMS ramp and when the quadrupole changes the mass position, the MS data is recorded simultaneously. However, these simultaneous events in synchro-PASEF bring their own challenges since there are several delays between triggering the programmed isolation position and recording the fragment spectra: First, the time constant of the electronics and thus the rise time of the actual values that are applied to the quadrupole rods need to be adapted to each other; Second, the ions’ mass and mobility-dependent transition time from the quadrupole to the TOF detector need to be accounted for. Both result in an apparent isolation mass shift of the quadrupole. Moreover, the transition time of ions through the quadrupole reduces the effective isolation width, given the very high scanning speed. To correct for this, we quantified the combined effects in terms of isolation mass and width offsets for scan rates up to 36,000 Th/s (see below) and determined linear correction functions to calibrate the quadrupole isolation mass and width in alignment with the synchronous movement.

### Experimental design and statistical rationale

We performed the experiments with tryptic HeLa cell lysate digest obtained from HeLa S3 cells (American Tissue Culture Collection, ATCC) to illustrate the behavior of synchro-PASEF in a complex sample type and with a simple tryptic protein digest mixture consisting of bovine serum albumin (BSA, P02769), yeast alcohol dehydrogenase (ADH, P00330), yeast enolase (P00924), and rabbit phosphorylase b (P00489) for detailed analyses. This simple protein mixture was obtained from Waters GmbH.

The complete dataset consisted of 13 raw files of single runs for this proof-of-principle study. For experiments involving a complex proteome mixture, 200 ng HeLa digest was used; for experiments involving the simple protein mixture, 200 fmol was injected for LC gradients of 3.2 min (300 samples per day - SPD on the Evosep system) and 5.6 min (200 SPD), and 400 fmol for LC gradients of 11 min (100 SPD) and 21 min (60 SPD). To analyze, simple protein digests in a complex background, we spiked 200 fmol protein mixture into 100 ng HeLa. The figure legends contain additional experimental design and statistical rationale for the respective experiments.

### Sample preparation

The complex proteome mixture used in our experiments was composed of HeLa S3 cells (ATCC). Cells were cultured in Dulbecco’s modified Eagle’s medium (Life Technologies Ltd., UK) supplemented with 20 mM glutamine, 10% fetal bovine serum, and 1% penicillin-streptomycin. The sample preparation was performed following the in-StageTip protocol (20). After washing the cells in PBS and cell lysis, the proteins were reduced, alkylated and digested by trypsin (Sigma-Aldrich) and LysC (WAKO) (1:100, enzyme/protein, w/w) in one step. The peptides were dried, resuspended in 0.1% TFA/2% ACN and 200 ng digest was loaded onto Evotips (Evosep, Odense, Denmark). The Evotips were prepared by activation with 1-propanol, washed with 0.1% formic acid (FA)/99.9% acetonitrile (ACN) and equilibrated with 0.1% FA. After loading the samples, tips were washed once with 0.1% FA.

The simple proteome mixture was composed in equal amounts of the purified and predigested tryptic standard proteins enolase, phosphorylase b, alcohol dehydrogenase (ADH), and bovine serum albumin (BSA). The peptides were reconstituted in 0.1% FA, and either 200 or 400 fmol (see above) were loaded onto Evotips.

### LC analysis

The Evosep One liquid chromatography system separated peptide mixtures at various throughputs using standardized gradients with 0.1% FA and 0.1% FA/99.9% ACN as the mobile phases. The 60 and 100 samples per day (SPD) runs were measured with an 8 cm × 150 μm reverse-phase column packed with 1.5 μm C_18_-beads (Bruker Daltonics) connected to a fused silica 10 μm ID emitter (Bruker Daltonics). For measuring 200 and 300 SPD, we connected the same emitter to a 3 cm × 150 μm reverse-phase column packed with 1.9 μm C_18_-beads (Bruker Daltonics). In both cases, the emitter was operated inside a nano-electrospray ion source (Captive spray source, Bruker Daltonics).

### MS analysis

The Evosep LC system was coupled with a modified timsTOF Pro mass spectrometer (Bruker Daltonics) to acquire data in synchro-PASEF mode and for comparison in dda- and dia-PASEF mode.

We acquired dda-PASEF with four PASEF scans per top-N acquisition cycle, an intensity threshold of 2,500 arbitrary units (a.u.) as indicated by the Bruker acquisition software, and a target value of 20,000 a.u. for precursor selection. Precursors that reached this target value were excluded for 0.4 min, which still permitted resequencing in simple mixtures, as shown in section two. Singly charged precursors were filtered out based on their m/z-ion mobility position, and precursors with a mass below 700 Da were isolated with a quadrupole selection window of 2 Th and otherwise with 3 Th.

For dia-PASEF, we used a method generated with our python tool py_diAID (16). That method covered an m/z-range from 350 to 1200 in twelve dia-PASEF scans with two isolation windows per dia-PASEF scan resulting in a cycle time of 1.4 s.

Additionally, we optimized three synchro-PASEF methods based on the tryptic HeLa reference library introduced in our optimal dia-PASEF study (16). The first method consists of four 25 Th isolation windows with a cycle time of 0.56 s and a second method of seven 15 Th windows (0.9 s cycle time). The third method had one 35 Th, two 25 Th, and one 50 Th isolation window (cycle time 0.56 s). The calculated precursor coverages and kernel density plots depicting a precursor ion cloud in the figures was based on this reference library, too.

If not stated otherwise, we used 100 ms to accumulate and elute ions in the TIMS tunnel. The m/z-range was acquired from 100 to 1700 and the ion mobility range from 1.3 to 0.7 Vs cm^−2^ calibrated with Agilent ESI Tuning Mix ions (m/z, 1/K_0_: 622.02, 0.98 Vs cm^−2^, 922.01, 1.19 Vs cm^−2^, 1221.99, 1.38 Vs cm^−2^). Experiments described to be ‘without collision energy’ were measured at a constant collision energy of 10 eV (because the software does not allow setting this value to zero), which is not sufficient to cause peptide fragmentation. For experiments with collision energy, the energy was linearly decreased from 59 eV at ion mobility of 1.6 Vs cm-^2^ to 20 eV at 0.6 Vs cm^−2^.

### Raw data analysis

Data was analyzed by manual and visual inspection with the help of our publicly available Python tools AlphaTims (21) and AlphaViz (22). The MaxQuant evidence output table (Version 1.6.17.0) (23, 24) from a single HeLa dda-PASEF run was the basis for calculating the percentage of precursor slicing. We used the reviewed human proteome (UniProt, Nov 2021, 20,360 entries without isoforms) and default settings for the analyses. This comprised a false discovery rate (FDR) of 1%, two missed cleavages, the cleavage pattern for trypsin, cysteine carbamidomethylation as fixed modification, and methionine oxidation and protein N-terminal acetylation as variable modifications. The mass tolerance was set to 10 ppm for the main search, and known contaminants were excluded from the dataset.

### Statistical analysis

The raw data was analyzed and visualized with Python (3.8, Jupyter notebook) and the packages pandas (1.2.4), AlphaTims (1.0.4) and numpy (1.20.3, (25)) for data accession and py_diAID (0.0.16), AlphaViz (1.1.15), matplotlib (3.4.2), and astropy (5.1) for visualization.

## RESULTS

### Synchronizing quadrupole movement with trapped ion mobility separation

The timsTOF principle adds a combined trapping and separation device to the regular liquid chromatography quadrupole time of flight (TOF) set up (26–28) (**Fig. 1A**). The TIMS tunnel has two segments, the first of which accumulates the ions for typically 100 ms, while the second releases the ions from the tunnel in high to low order, with the ions that have the largest cross sections (and typically mass) first. Although the TIMS device is capable of an ion mobility resolution of greater than 100, in complex mixtures the ion mobility peaks are typically several ms wide.

**Figure 1:**
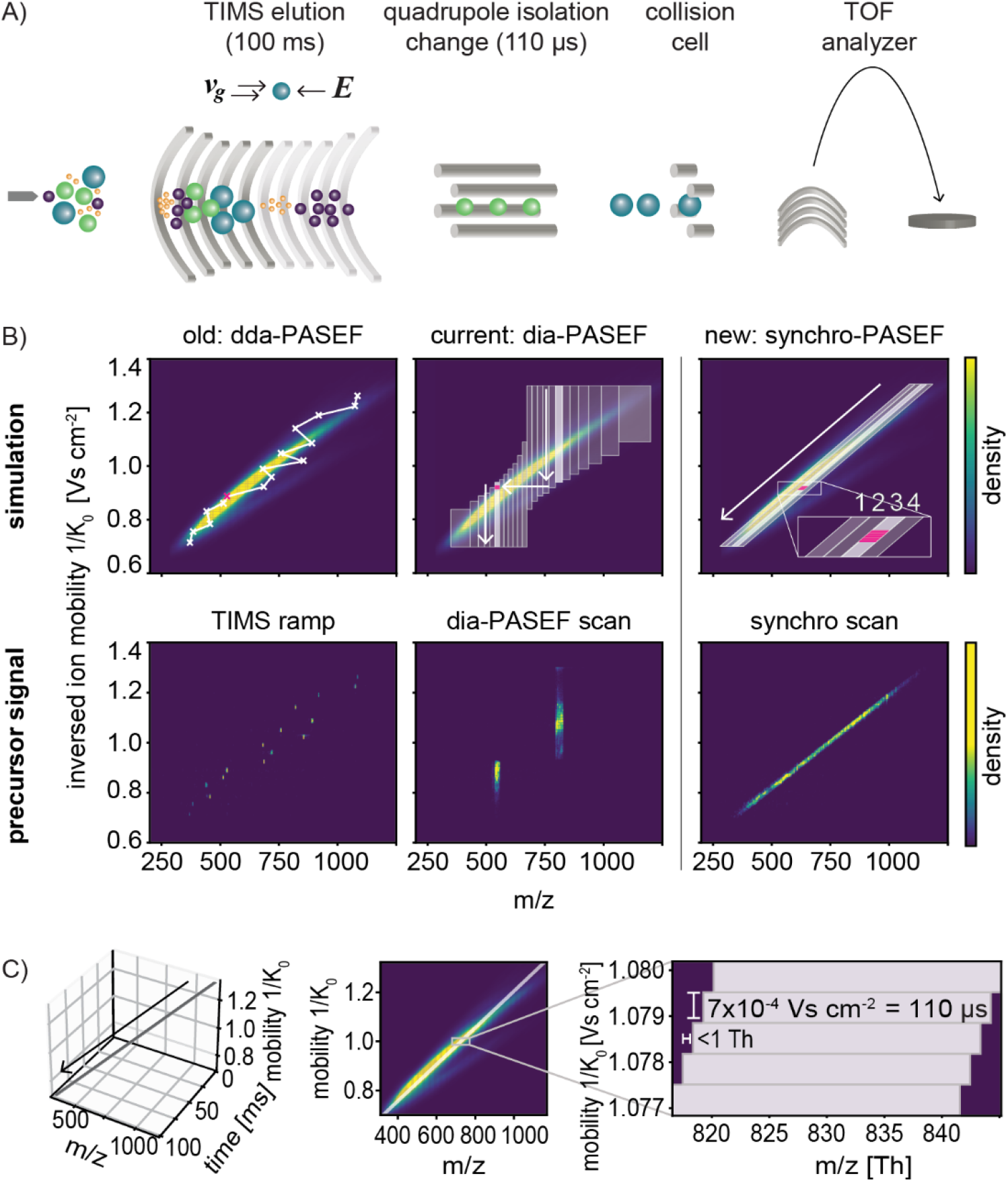
The principle of synchro-PASEF precursor isolation. A) The timsTOF ion path: Ions enter the mass spectrometer after electrospray, accumulate and separate in the TIMS tunnel, and are then isolated by a mass-selective quadrupole. A collision cell fragments these precursor ions and a reflector TOF analyzer determines the mass spectrum every 110 us. B) Top panel: simulations of dda-PASEF, dia-PASEF, and synchro-PASEF. The background ion cloud in m/z and ion mobility dimensions shows a kernel density distribution of a reference library from tryptic HeLa (see Experimental Procedures). Bottom panel: precursor signal of each scan mode from 200ng tryptic HeLa digest acquired without collision energy for a 21 min gradient (60 samples per day). dda-PASEF selects the top-N most intense precursors with an isolation width of 2-3 Th. Each TIMS ramp isolates different precursors, and the 5656^th^ TIMS ramp is visualized above (left panel). dia-PASEF isolates the precursor cloud in vertical stripes with variable isolation widths. A 20 Th isolation in the 6^th^ dia-PASEF scan at frame 5649 is shown (middle panel). Synchro-PASEF follows the precursor cloud naturally and selects many more precursors per isolation window as the previous dia-PASEF schemes. The 3^rd^ synchro scan, frame 5654, is shown (right panel). C) Left panel: Visualization of the synchro-PASEF quadrupole movement in m/z, inverse ion mobility and TIMS ramp time dimensions. The quadrupole control steps 0.9 Th to lower m/z values for every 110 μs TOF spectrum, corresponding to a 0.0007 V cm^−2^ downwards step in the inverted ion mobility direction.

The downstream quadrupole sets the molecular mass per charge (m/z or Th) isolation range of interest, and the selected precursors are subjected to fragmentation, with all fragment ions appearing at the same ion mobility value as their precursors. The PASEF principle utilizes the speed of the TOF analyzer and the correlation between ion mobility and molecular weight to subject more than one precursor per accumulation cycle for fragmentation. This implies a selection in two dimensions since all peptide precursors require isolation based on molecular weight and ion mobility (13, 15). Note that the selection in m/z dimension is performed explicitly by the quadrupole whereas the selection in ion mobility dimension happens implicitly by the timing of the quadrupole selection in relation to the release of ions from the TIMS device.

In DDA mode, PASEF allows the selection of more than ten precursors per TIMS ramp, which involves rapid placement of the selection window to the precursor positions in a zig-zag pattern (**Fig. 1B**, left panel). In our previous implementation of DIA, the quadrupole is instead stationary with an appropriate selection window for a part of the TIMS ramp and then jumps the next mass range (**Fig. 1B**, middle panel). Although highly efficient (16), neither scan mode naturally follows the ‘banana-shaped’ ion cloud in the m/z vs. ion mobility plane.

We reasoned that a continuous and synchronized movement of the quadrupole could slice the ion cloud into several, equally complex TIMS ramps (**Fig. 1B**, right panel). To enable such a synchro-PASEF scan, we built upon the field programmable gate array (FPGA) already present in the instrument that previously controlled among others the TIMS scan. In addition, the digital analog converters (DACs) of the quadrupole RF-amplitude, the bias-DC, and the resolution-DC are all routed through this FPGA (Experimental Procedures). This allows the quadrupole to move continuously and extremely fast, covering about 8,000 Th/s while being aligned with the TIMS release (**Fig. 1C**).

In this particular implementation, the mass spectrometer executes 927 TOF pulses during the 100 ms TIMS ramp, where each TOF acquisition takes 110 μs. The leading edge (lower m/z) of the quadrupole selection window of 25 Th starts at m/z 1170 and moves down to 335 in our method (**Fig. 1C**). Thus, the quadrupole control changes its position every 0.11 ms with a step size of 0.9 Th. In comparison, Scanning SWATH (19) moves the selection window every 2 ms with a step size of 2 Th. Given a width of the ion cloud of about 100 Th of the doubly charged precursors, we found that four TIMS ramps with our 25 Th isolation width adequately cover these precursors. Based on the reference library, this leads to a precursor coverage of 76% (see Experimental Procedures). Adding one MS1 level TIMS ramp to measure precursors and transfer time, the total cycle time is only 0.56 s, much faster than our previously optimized dia-PASEF method for short gradients (1.4s, Experimental Procedures). Those methods were designed for comprehensive coverage, but only contain 8% of precursors per isolation window, in contrast to 19% for synchro-PASEF. The underlying reason for this advantage is that slicing the banana-shaped ion cloud vertically as we had done before requires many more steps than slicing it diagonally along its length while keeping the same width. Thus synchro-PASEF is better adapted to the ion cloud, thereby increasing precursor sampling and making it very fast and efficient.

### Synchro-PASEF combines continuous fragmentation with short cycle times

Having established the feasibility and parameters of synchro-PASEF scans on unfragmented precursor selection, we next investigated their benefits for the fragment spectra compared to dda- and dia-PASEF.

As mentioned above, dda-PASEF selects single precursors leading to a striped pattern of fragment ions in the ion mobility (**Fig. 2A**, left panel). Only a small proportion of precursors is covered in each TIMS ramp, but for these the precursor-fragment and the fragment-fragment relationships are clearly defined. These fragmentation spectra can still be composed of more than one co-eluting precursor in the same precursor isolation window, but the ion mobility selection makes this less likely.

**Figure 2:**
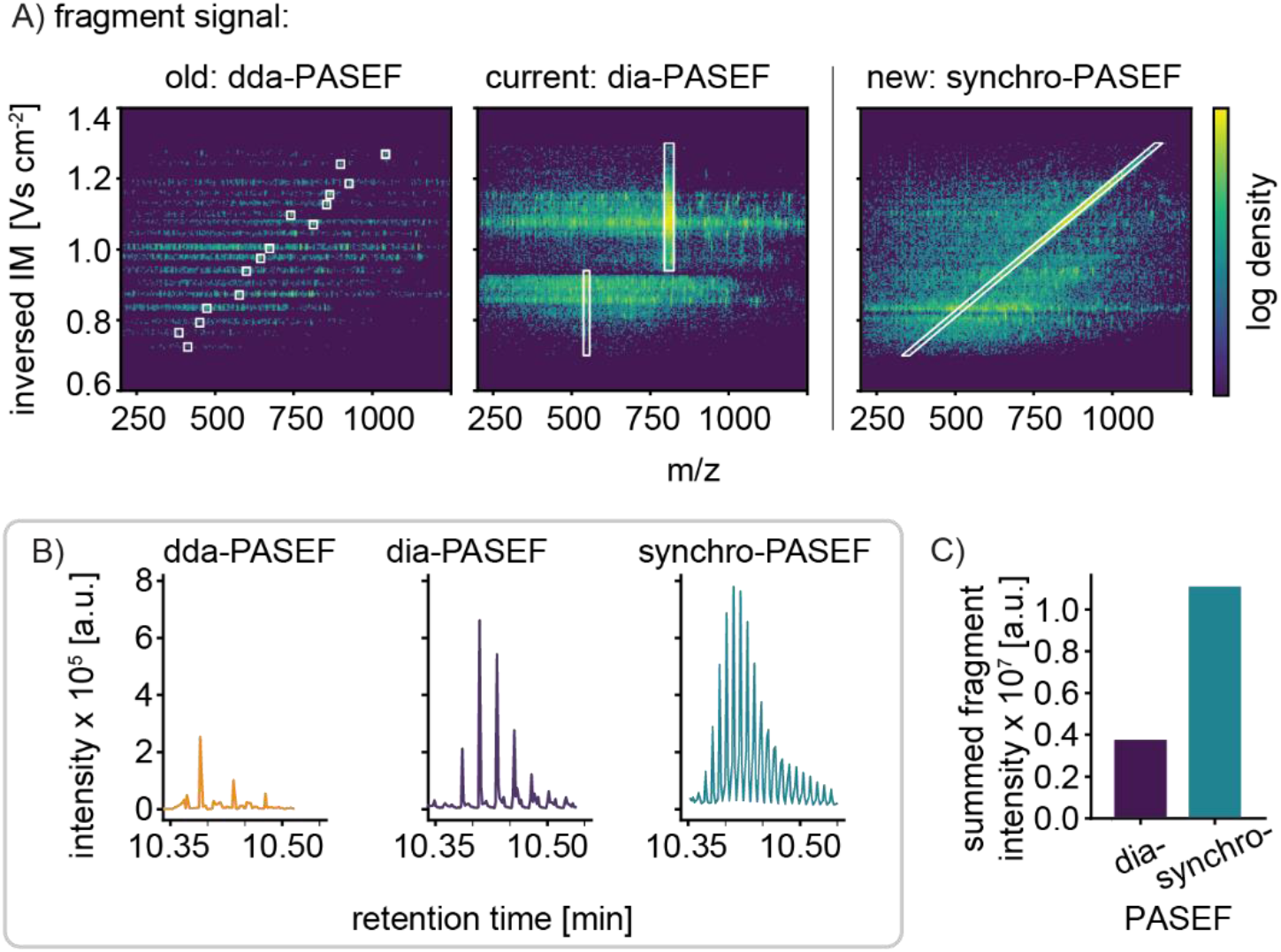
Fragmentation behavior for dda-, dia- and synchro-PASEF. A) Comparing fragment signals of dda-, dia- and synchro-PASEF. A tryptic digest of HeLa cells was measured in a 21 min gradient. The same frame numbers as in Figure 1B are shown but collision energy was applied. Left panel: fragmentation spectra of 15 precursors marked with their isolation windows in dda-PASEF. Middle panel: isolation windows and continuous fragment acquisition in dia-PASEF. There are no fragments at the transition point and at the precursor low mobility regions. Right panel: The isolation window of the third synchro-PASEF scan is shown. Note the uniform fragment distribution in the entire m/z – ion mobility plane. B) Conversion efficiency to fragments for the three scan modes illustrated with the fragment intensities of GLAGVENVTELK (phosphorylase b) and SISIVGSYVGNR (ADH) co-eluting at a retention time of 10.35 to 10.55 min and an ion mobility of 0.93 to 0.955 Vs cm^−2^. In DDA mode the eluting peptide was fragmented three times in this low complexity mixture. Synchro-PASEF sampled the peptide many more times than dia-PASEF due to shorter cycle times. C) Summed intensities for the fragment signals of the two peptides in (B).

High data completeness is inherent to DIA modes since a broad mass range is recorded deterministically. Furthermore, DIA samples the precursor and fragments multiple times over an elution peak in the retention time and – in the case of dia-PASEF – in the ion mobility dimension. This sampling over the elution peaks in principle enables the reconstruction of the relationship between the precursor and its fragments based on the alignment of the elution profiles. However, the broad isolation windows of DIA lead to highly chimeric fragment spectra since multiple precursors are selected at once, which in practice limits a precise reconstruction of these relationships even in dia-PASEF (**Fig. 2A**, middle panel).

Synchro-PASEF maintains the principle of DIA - fragmenting multiple precursors at once – resulting in complex spectra. However, in contrast to our previous dia-PASEF methods, the fragmentation spectrum is continuous in the m/z and ion mobility dimension, filling nearly the entire m/z vs. ion mobility plane. Furthermore, there are no artifactual borders due to the quadrupole jumping to different mass ranges (**Fig. 2A**, right panel).

To directly measure the increase in fragmentation yield in synchro-PASEF, we analyzed a low complexity mixture (Experimental Procedures). Taking the co-eluting phosphorylase b peptide GLAGVENVTELK and ADH peptide SISIVGSYVGNR measured with 21 minutes gradients (60 SPD on the Evosep system) as an example, we only observed a few fragmentation events across the elution peak in DDA (caused by ‘resequencing events’ in this low complexity mixture) (**Fig. 2B**, left panel). In contrast, our recently reported optimal dia-PASEF method (16) had five fragmentation data points per peak at a cycle time of 1.4 s (**Fig. 2B**, middle panel). As outlined above, the diagonal quadrupole movement of synchro-PASEF leads to much more rapid sampling of the eluting peaks, in this case multiplying the number of fragmentation data points compared to dia-PASEF (**Fig. 2B**, right panel). Importantly, the increased sampling also amplifies the integrated signal for each fragment. For these peptides, synchro-PASEF acquired three-fold more fragment signal than our previous dia-PASEF method, as expected from the nearly three-fold faster sampling (**Fig. 2C**).

### Extensions of synchro-PASEF

So far, we have elucidated the features of synchro-PASEF with an acquisition method that had a constant isolation width of 25 Th, four synchro scans, a ramp time of 100 ms and a quadrupole step size of 0.9 Th. This is, however, just a reference method and can be adjusted depending on the use case, for example to increase proteome coverage. Next, we explored three acquisition schemes that achieve different balances between quadrupole isolation width, cycle time, precursor coverage, and now also quadrupole step size (**Fig. 3A**).

**Figure 3:**
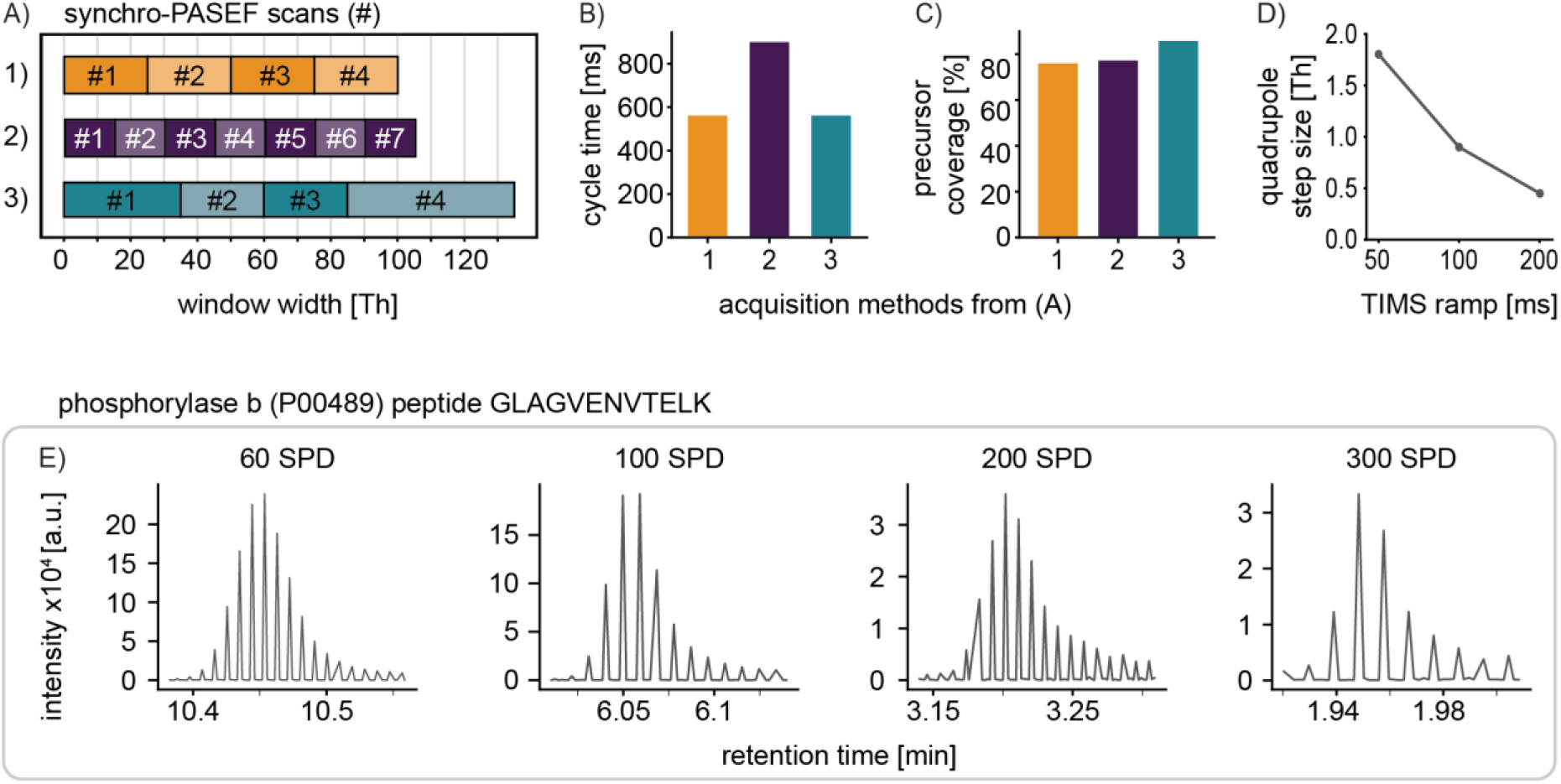
Parametrization of synchro-PASEF scan modes. A) Acquisition schemes for synchro scan methods with different quadrupole selection window widths. B) Cycle times of the methods in (A). C) Precursor coverage of methods in (A) calculated with the HeLa reference library. D) Quadrupole step size in dependence on the TIMS ramp time. E) Precursor elution profiles and sampled data points in the retention time dimension at a throughput of 60 SPD (21 min), 100 (11 min), 200 (5.6 min) and 300 (3.2 min) for the peptide GLAGVENVTELK.

The first one is the method predominantly used in this paper, the second one has seven 15 Th synchro scans, and the third method has four synchro scans with different window sizes adapted to the precursor cloud density but kept constant throughout each individual synchro scan. Since the second method has seven synchro scans, it also has the longest cycle time (900 ms instead of 561 ms, **Fig. 3B**). Method three has the widest isolation windows and covers 86% of the precursor cloud, which is 10% more than the coverage of method one and two, while having a short cycle time (**Fig. 3C**).

Given the constant number of TOF triggers (927 for 100 ms), the TIMS ramp time directly determines the quadrupole step size (when scanning the same m/z range). At a ramp time of 100 ms, the TOF executes 927 triggers in the standard methods, whereas doubling the ramp time also doubles the number of TOF triggers to 1854. In the case of a quadrupole selection window of 25 Th moving from 1170 to 335 Th, a 200 ms ramp time leads to a step size of 0.45 Th and 50 ms to 1.8 Th (**Fig. 3D**). Thus, the quadrupole step size can be made even smaller for longer timsTOF ramps.

Independent of the ramp time, the quadrupole changes position every 110 μs (the duration of one TOF acquisition). A TIMS ramp 50 ms implies a step size of 1.8 Th and a quadrupole scanning speed of about 16,000 Th/s. We characterized the isolation performance at these very fast scan rates to evaluate the implications for method optimization. Note that the electronics trigger the programmed mass position earlier than the actual recording of the fragment spectrum (Experimental Procedures). We quantified the isolation mass and width effects for scan rates up to 36,000 Th/s and determined linear calibration functions to correct for the synchronously moving quadrupole (**supplementary Fig. 1A**). After calibration, the mass deviation of the center of the 25 Th isolation window at a scan speed of 16,000 Th/s Th/s was less than ± 0.4 Th for the relevant mass-to-charge range. Deviation of total isolation width was less than ± 0.5 Th (**supplementary Fig. 1B**). Note that this is not the accuracy of the leading and trailing edges of the isolation window (see below).

Motivated by this quadrupole precision even at very high scan rates, we wished to investigate how much LC gradients could be shortened while retaining many data points per peak. We acquired our simple protein mixture with the standard method (four times 25 Th synchro scans) with progressively higher throughput (**Fig. 3E**). Taking the same peptide GLAGVENVTELK for illustration, we achieve high peak sampling for all gradients, including the ones with FWHM peak widths around one second. Remarkably, even at a throughput of 300 samples per day (SPD) corresponding to an LC gradient of 3.2 minutes, we still obtained eight data points per peak with a cycle time of 0.56 ms. This suggests that synchro-PASEF is suitable for high throughput, quantitative DIA applications.

### Precursor slicing by synchro-PASEF deconvolutes DIA spectra

Our results up to this point establish superior precursor sampling efficiency and speed for synchro-PASEF compared to previous scanning modes. However, when analyzing the data we noticed that the diagonal synchro scans frequently intersected or ‘sliced’ precursors in the m/z and ion mobility plane. In fact, this was the rule rather than the exception and is caused by the fact that scan lines are closer together than the ion mobility peak widths (see below). We also noticed that precursor slicing divides the fragment signals into adjacent synchro scans.

This principle is depicted schematically in Figure 4A. When the quadrupole moves from high to low m/z values and from high to low inversed ion mobility values, the trailing edge of the quadrupole selection window (synchro scan #1) moves through the orange precursor and the top parts of all fragments for this precursor are recorded. The leading edge of the next synchro scan slices the precursor at the same position, this time recording the bottom parts of all fragments. Importantly, adding the top and bottom parts reconstitutes the total fragment signal with the same two-dimensional peak shape as the precursor signal.

**Figure 4:**
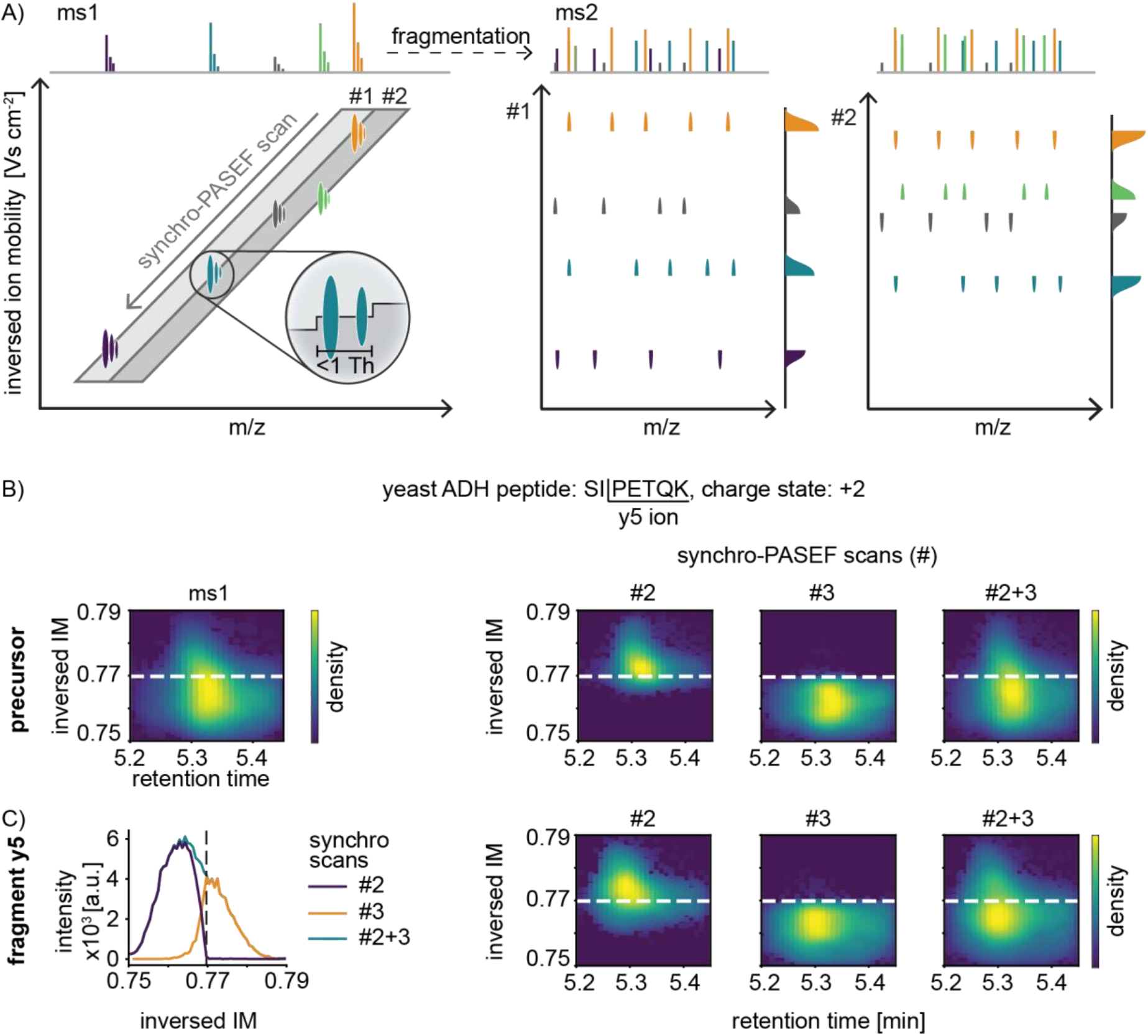
Principle of precursor slicing in synchro-PASEF. A) Two synchro scans and five precursors are schematically depicted (left panel). As the trailing edge of synchro scan #1 moves through the orange precursor, the top parts of all fragments appear at the same time, followed by the bottom part in scan #2 (middle and right panels, respectively). The slicing position is known for each precursor, and all pieces with the same slicing position belong together. This specifies the precursor-fragment and fragment-fragment relationships. B) Slicing of the quadrupole of the yeast ADH peptide SIPETQK. For illustration, no collision energy was applied. The precursor peak can be quantitatively reconstructed from the sliced parts in two subsequent synchro scans. C) Same as (B) but applying collision energy and focusing on the y5 ion. The traces in adjacent synchro scans add up to the full trace for this transition (left panel). The fragment is sliced in exactly the same way as the precursor in (B) (right three panels).

To analyze how frequent such slicing events occur, we note that at each quadrupole position the isolation window is 25 Th wide, and the scan diagonal of one synchro-PASEF scan is 3.2 ms high in the ion mobility dimension (**supplementary Fig. 2A**). With our dda-PASEF HeLa run from Figure 2A, we determined that the average ion mobility peak width was 4.3 ms (**supplementary Fig. 2B**). A total of 79% of the precursor peaks have an ion mobility profile wider than the synchro scan height. Altogether, 97.4% of the precursors that are within the area covered by the synchro scans are sliced at least once, making precursor slicing a nearly universal phenomenon. The percentage of precursors that are sliced depends on the height of the synchro scans in ion mobility direction, which in turn varies with the quadrupole selection window width and the TIMS ramp time. Referring back to the three synchro-PASEF methods in Figure 3A, the wider isolation windows of method 3 reduce the percentage of sliced precursors to 90.1%. Method 2 with the smallest isolation width (15 Th) also has the smallest height of the diagonal, resulting in 99.7% of precursors being sliced (**supplementary Fig. 2C**).

To experimentally demonstrate precursor slicing, we measured our simple protein mixture, focusing in detail on the doubly charged yeast alcohol dehydrogenase peptide SIPETQK (modified sequence: acetylated protein N-term-SIPETQK, m/z 422.7). Acquiring this peptide without collision energy in MS1 mode defines the peak shape in retention time and ion mobility space (**Fig. 4B**, leftmost panel). Given the known precursor m/z and ion mobility it is straightforward to calculate the intersection point of the synchro scan that slices this precursor (dashed line in the figure). Applying synchro scans without collision energy, the top part of the precursor appears in one synchro scan with the edge at the expected slicing position. The next synchro scan contains the remainder of the peak and both parts together reconstitute the full precursor peak (**Fig. 4B**, rightmost panel). This peak reconstitution can also be seen when signal intensities of the two synchro scans are separately plotted as a function of m/z, retention time or ion mobility (**supplementary Fig. 3A**).

Next, we examined how precisely the quadrupole slices. Inspection of the ion cloud after synchro scan isolation but without fragmentation showed good overall alignment between the programmed isolation window and actually transmitted ions, with the leading edge of the quadrupole ejecting lower mass ions somewhat sharper than the trailing edge ejecting higher mass ions (**supplementary Fig. S3B, C**). This is the expected behavior because the stability to instability transition of a quadrupole is steeper for the low mass ejection than the one for high mass ejection (29). When applying collision energy, we observe the slicing behavior for all fragments (shown for y5 in **Fig. 4C**).

Synchro-PASEF inherits the PASEF dimensions retention time, ion mobility, m/z, and intensity from dia-PASEF. However, precursor-fragment or fragment-fragment relationships are defined by the phenomenon of precursor slicing. As illustrated in our simple mixture and in more detail below, two-dimensional precursor slicing conceptually ‘deconvolutes’ the complex DIA spectra because it establishes precursor-fragment relationships similar to DDA spectra. In the next section, we refine this concept and apply it to qualitatively and quantitatively disentangle individual peptides from the complex background of a cellular proteome.

### Precise identification and quantification via a lock and key mechanism

As described above, synchro-PASEF slices precursors and forms complementary fragment parts in adjacent synchro scans. When put back together at the known slicing position of the precursor, these parts should reconstitute its shape. We found these criteria to be exquisitely specific (see below), reminiscent of a ‘lock and key mechanism’.

To illustrate, we first depict a precursor and the slicing plane in the m/z, retention time and ion mobility dimensions (**Fig. 5A**). A number of fragment ions have roughly similar retention times and ion mobilities and could therefore potentially belong to the precursor (**Fig. 5B**). From the calculated slicing position of the precursor, the potential fragments can be verified based on their experimental slicing pattern (**Fig. 5C**). Investigating these peaks more closely, fragments 1, 3, and 4 have the same shape and slicing position as the precursor and are correct identifications. Fragment 2 co-elutes in retention time and ion mobility but is sliced at a different position then calculated for the precursor mass. Fragment 5 has a different shape in the ion mobility dimension and therefore cannot be fragments for this precursor.

**Figure 5:**
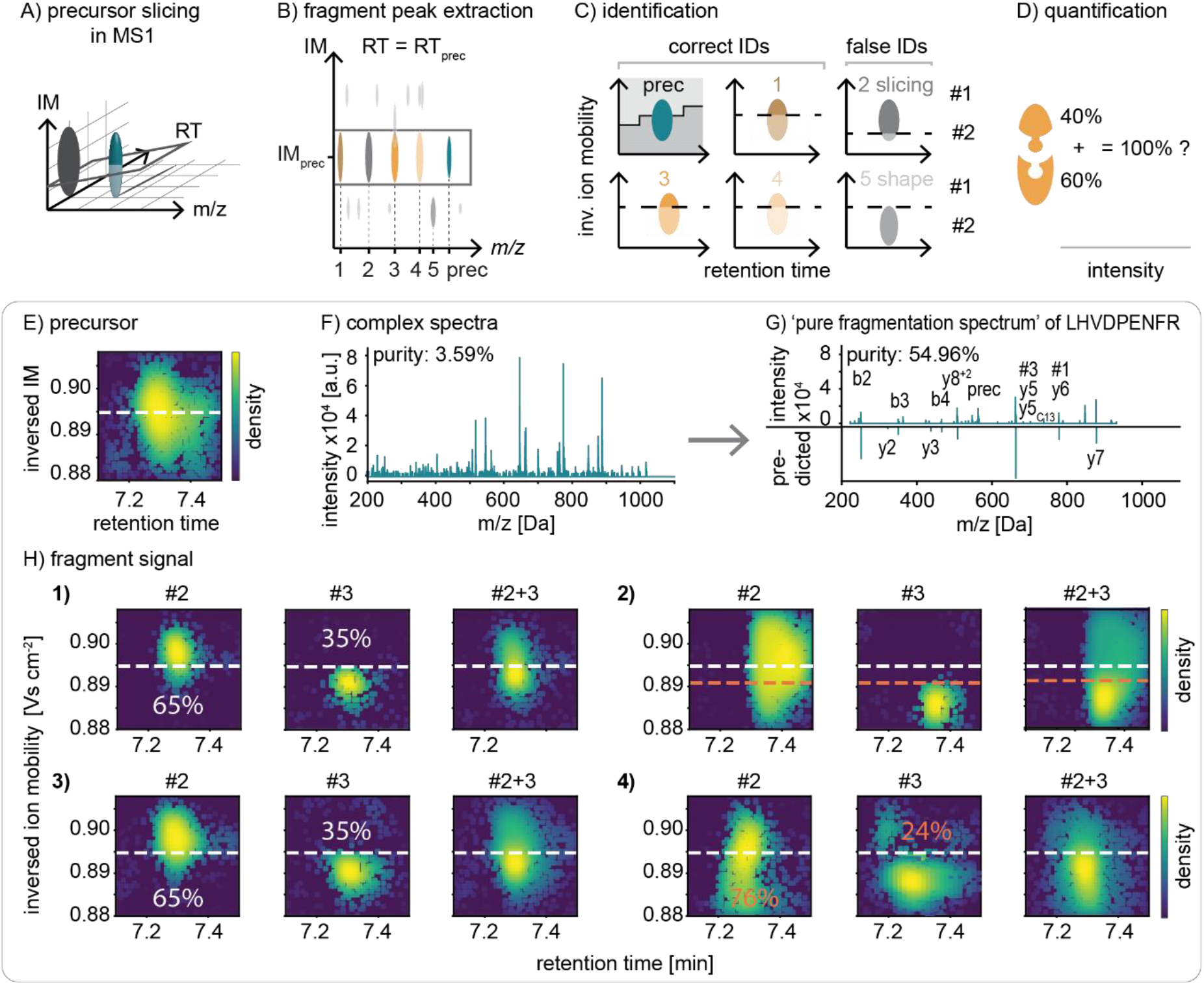
Precursor slicing makes identification and quantification more precise. A) The quadrupole slices precursors into at least two parts. B) All features with the same retention time and ion mobility distribution as the precursor are potential fragments of the precursor. C) Precise identification enabled by slicing: Fragment 2 is sliced at the wrong position, and fragment 5 does not align in the retention time and ion mobility dimension; hence these fragments do not belong to this precursor. D) Slicing improves the confidence for correct quantification since both sliced parts should add up to 100% normalized intensity. E) A particular precursor in the HeLa digest and its calculated slicing position indicated by a dashed line. F) Fragment m/z spectrum extracted at the precursor’s ion mobility and retention time position. G) Pure fragmentation spectrum after inspecting the single fragments for correct precursor slicing and the mirrored spectrum predicted by AlphaPeptDeep (6). H) Fragment signal from (F) in detail.

Precursor slicing has also implications for quantification based on the requirement that fragment parts added together should reconstitute the shape of the precursor peak. The fraction of the precursor intensity in each synchro scan have fixed ratios given by the slicing position (**Fig. 5D**). If the top and bottom parts of a sliced fragment do not add up to the intensity distribution of the precursor, this typically indicates an interference of another precursor at this fragment mass.

We demonstrate the specificity of this principle on a precursor, which we calculate to be sliced at an ion mobility position of 0.895 Vs cm^−2^ based on a precursor m/z of 563.786 (**Fig. 5E**). The fragmentation spectrum for this precursor is complex even after filtering for concordance in retention time and ion mobility (**Fig. 5F**) but this can be disentangled by inspecting the individual peaks in detail (**Fig. 5G**). The prominent peaks labeled signals 1 and 3 have a similar shape and slicing position as the precursor. Additionally, their intensity ratios (35% and 65%) correspond to the calculated precursor slicing (**Fig. 5H**). Fragment 2 is sliced at a different position and has a different shape. Fragment 4 has the correct slicing position, but the fragment part from synchro scan two is overlayed with another signal, changing the intensity ratios. As a result, only fragments 1 and 3 are valid. Note that only one part of fragment 4 has an interference, therefore the second part could still be used for identification and quantification.

Going through all fragment signals leads to an extracted spectrum as simple or even simpler than a DDA spectrum (**Fig. 5G**). The precursor and fragment masses extracted in this way excellently fit the doubly charged peptide LHVDPENFR (P02081) of hemoglobin fetal subunit beta that is present in the HeLa growth medium. Fragment 1 is the y6 ion, fragment 3 the y5, and fragment 4 the y7 ion. Going through all potential fragment peaks of this precursor yields an extremely clean fragmentation spectrum, nearly devoid of unassigned fragments (see explanation in supplementary Figure 4). Moreover, precursor slicing also allows the assignment of peaks other than backbone fragmentation that would not normally be assigned by a search engine.

To demonstrate the extraction of DDA like spectra more objectively, we assigned precursors of the simple protein mixture that had been spiked into a complex HeLa mixture. We then filtered for fragments belonging to each precursor as described above, except that we defined a correlation threshold of at least 0.3 and deviation from the intensity ratios of less than 3 for retained fragments. Because virtually all fragments in the extracted spectra belong to one precursor (unlike DDA spectra), we term them ‘pure fragmentation spectra’. The ion current in the 29 pure spectra that is easily explainable by the expected fragmentation spectrum is 55.6% on average, whereas this was 4.6% without extraction (**supplementary Fig. 4**).

## DISCUSSION

In this paper, we have developed and characterized a novel scanning method termed synchro-PASEF. A defining feature is that the quadrupole selection window moves diagonally through the m/z - ion mobility plane and synchronously to the ion mobility elution. In this sense, it is the two-dimensional extension of methods such as SONAR or Scanning SWATH. Compared to our previous DDA or DIA based PASEF methods, this movement samples the precursor cloud more efficiently, covering up to 19% of all precursors per isolation window. Additionally, it allows short cycle times which is reflected in up to three-fold higher fragment intensities and much higher peak sampling rates compared to our previous standard method. This implies adequate sampling even for very short LC gradients, in turn opening up for high throughput. The fragment signal amplification and higher sampling rate should also be beneficial for quantitative precision.

Another key and unanticipated feature of synchro-PASEF is the fact that precursors are sliced in m/z and ion mobility space by the fast-moving quadrupole selection window. Nearly all precursors are sliced and their fragments are thereby separated into adjacent synchro scans with high precision and reproducibility. The slicing position deterministically depends on the precursor mass and establishes precursor-fragment and fragment-fragment relationships. We found that the separated fragment parts can be recombined quantitatively with very high specificity. This lock and key mechanism can serve as a highly specific criterium for identification and quantification, in effect deconvoluting the complex DIA spectra to the simplicity of DDA spectra or perhaps even better. We plan to use this additional information to eliminate false identifications, pinpoint signals with interference and even correct them by retrieving only the interference-free part of the transition.

Taken together, synchro-PASEF adds another dimension to the already five-dimensional data cube in timsTOF - consisting of retention time, ion mobility, quadrupole selection, m/z, and intensity (28). It offers all the advantages of data-independent acquisition, including a comprehensive picture of precursor and fragments over the elution profiles, as well as high data completeness resulting from a high ion beam usage with short cycle times. This is combined with the specificity of DDA owing to a defined precursor-fragment and fragment-fragment relationship established by two-dimensional precursor slicing. Clearly, both library-free and library-based analysis could benefit from this principle. In this sense, synchro-PASEF promises to combine the advantages of DDA and DIA.

Here, we only established these defining features and potential of synchro-PASEF but did not actually evaluate proteome coverage or quantitative accuracy in complex proteome mixtures at a global scale. This is due to the current absence of tools that can handle the data structures generated by the synchro scans. Once the appropriate algorithms and software are available, it will be interesting to see how the principal advantages of synchro-PASEF translate to increases in proteome depth, coverage and quantitation. This will also be a good point to optimize and extend the method parameters, similar to what we reported for dia-PASEF to design optimal methods (16). For instance, synchro scan borders could be chosen somewhat wider (as in method 3 in Figure 3A) to cover more of the precursors. Likewise, the borders of the quadrupole selection windows could be offset between adjacent scans to ensure that all precursors are sliced.

Once the downstream informatic infrastructure is in place, synchro-PASEF may be particularly promising for proteome types that are traditionally challenging for DIA. We speculate that the additional confidence in fragment identity would be useful for studying post-translational modifications, where each additional fragment can be crucial in pinpointing the modification site. Likewise, synchro-PASEF may help disentangle the very complex spectra generated by multiplex DIA approaches (30–32). Since precursor slicing is directly dependent on the precursor mass it could link fragments to labeled precursors that are co-eluting and are co-isolated for fragmentation.

## ABBREVIATIONS

ADH: alcohol dehydrogenase
BSA: bovine serum albumin
DAC: digital analog converter
DC: direct current
DDA: data-dependent acquisition
DIA: data-independent acquisition
FA: formic acid
FPGA: field programmable gate array
IM: ion mobility
PASEF: parallel accumulation – serial fragmentation
RF: radio frequency
RT: retention time
SPD: samples per day
TIMS: trapped ion mobility spectrometry

## DATA AVAILABILITY

Homo sapiens (taxon identifier: 9606) proteome database and the sequence of BSA (P02769), ADH (P00330), enolase (P00924), phosphorylase b (P00489) were downloaded from https://www.uniprot.org.

## SUPPLEMENTAL DATA

This article contains supplemental data.

## ACKNOWLEDGEMENT

We thank our colleagues at the department of proteomics and signal transduction as well as at Bruker Daltonics for discussions and help. We are grateful for the input from Igor Paron, Bianca Splettstoesser and Fabian Opitz. Maximilian Strauss suggested using the cross correlation for retrieving fragments. This study was supported by the Max-Planck Society for Advancement of Science, the Deutsche Forschungsgemeinschaft project “Chemical proteomics inside us” (grant no.: 412136960), European Union’s Horizon 2020 research and innovation program under grant agreement No 874839 ISLET and by the Bavarian State Ministry of Health and Care through the research project DigiMed Bayern (www.digimed-bayern.de).

## AUTHOR CONTRIBUTIONS

P.S., S.W. O.R. and M.M. conceptualized and designed the study; F.K., M.L., and O.R. developed the technological innovations on the timsTOF, P.S., G.W., M.W., E.I., P.K., and F.M. and performed experiments; P.S., F.K., G.W., P.K., M.T., F.M., S.W. and M.M. analyzed the data; P.S. and M.M. wrote the manuscript with input from all authors.

