## supplementary figures 1 to 4 for "Synchro-PASEF allows precursor-specific fragment ion extraction and interference removal in data-independent acquisition"

### Supplementary figures 1 to 4 of:

### Synchro-PASEF allows precursor-specific fragment ion extraction and interference removal in data-independent acquisition

Patricia Skowronek, Florian Krohs, Markus Lubeck, Georg Wallmann, Ericka Itang, Polina Koval, Maria Wahle, Marvin Thielert, Florian Meier, Sander Willems, Oliver Raether, Matthias Mann

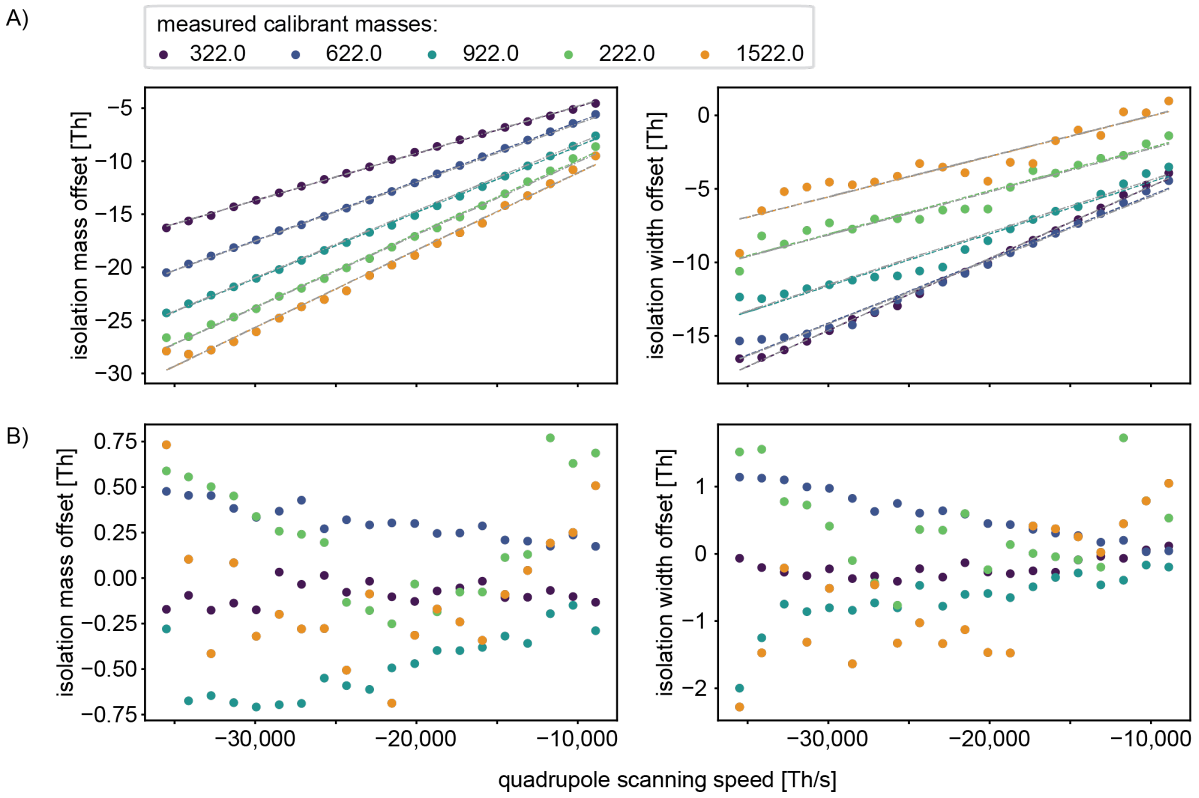


**Supplementary Figure 1: Precision of the quadrupole depending on quadrupole scanning speed.**

For this experiment a constant quadrupole isolation width of 25 Th was used.

A) Before quadrupole calibration. The position of the isolation window shows linear behavior with the quadrupole scanning speed (left panel). The width of the isolation window behaves less linear than the position when varying the quadrupole scanning speed (right panel)

B) After quadrupole calibration. The resulting isolation mass position offset is within ± 0.75 Th (left) and the resulting offset of the isolation window width is about ± 2 Th.

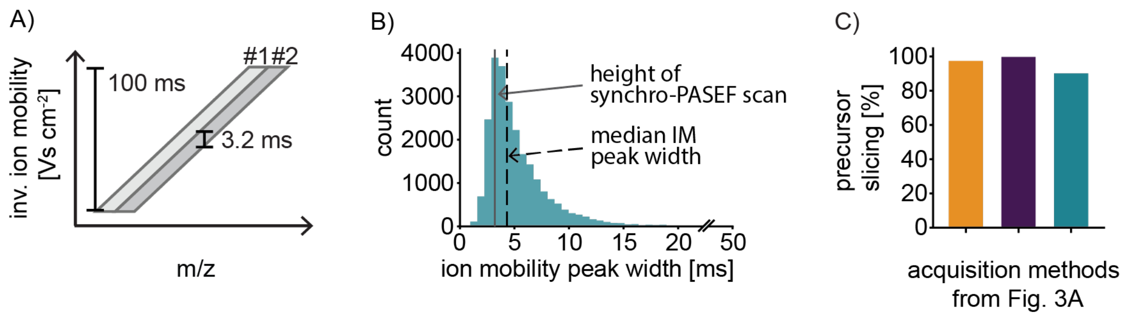


**Supplementary Figure S2: Precursor slicing is a nearly universal phenomenon.**

A) The TIMS tunnel releases precursor ions according to their ion mobility for 100 ms. The diagonal scan line has an isolation width of 25 Th and a height of 0.019 Vs $cm^{-2}$, corresponding to 3.2 ms.

B) Histogram of ion mobility peak widths from the HeLa reference library (see Experimental Procedures). The median is at 0.026 Vs $cm^{-2}$, corresponding to 4.3 ms. 21% of the peaks have an ion mobility width smaller than the height of the diagonal scan line.

C) Percentage of precursor slicing calculated with the dda-PASEF, HeLa run from Figure 2A. Note that only precursors with ion mobility within the four scan lines are considered.

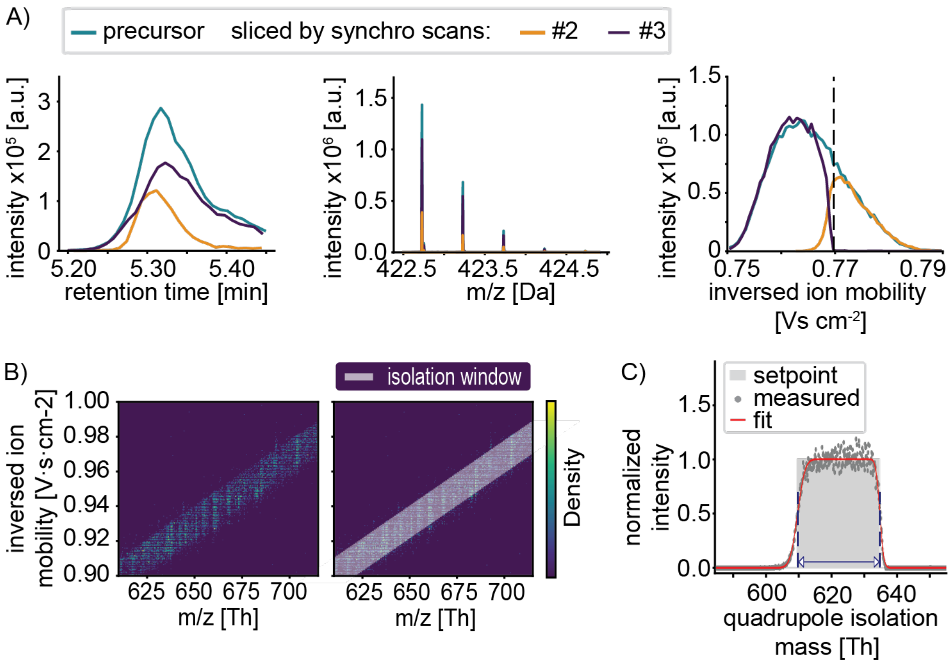


**Supplementary Figure S3: Precursor slicing in the ion mobility dimension**

D) The plots show the peptide SIPETQK of the simple protein mixture acquired without collision energy. The precursor signal and the signals, that are sliced by the synchro scans, align in the retention time dimension (left panel). The isotopic peak patterns of the adjacent synchro scans follow the expected intensity pattern (middle panel). The ion mobility dimension shows precursor slicing and confirms that both sliced peak parts conform to the precursor peak shape (right panel).

E) The left panel shows the isolated signal of a single synchro scan. In the right panel, the isolated signal is overlayed with the programmed isolation window in grey. The heatmaps demonstrate that the leading edge is exact and the trailing edge is somewhat softer.

F) The programmed quadrupole shape is rectangular (colored in light grey) and the actual quadrupole shape in red is fitted based on measured data points.

**Supplementary Figure 4 description: 'pure fragmentation spectra' from complex mixtures**

For a preliminary but unbiased evaluation of the discriminatory power of precursor slicing, we analyzed the simple protein mixture spiked into a complex cellular lysate background (see main text). A data set was generated consisting of the 268 peptides identified by a MaxQuant search (1) from which the precursor position and dimensions were retrieved. Fragment intensities were predicted in silico using AlphaPeptDeep (2) for comparison in mirrored spectra.
We set out to generate pure fragmentation spectra for the 40 peptides around the dataset's median intensity value for the 268 peptides. While all precursors were sliced, only the 29 of them that had at least two recognized parts were considered for our analysis.
The peptide fragment extraction window was positioned based on the peptide's retention time, ion mobility, and m/z ratio and recentered to the mass based on the intensity, only considering intensities with at least 25% of the maximum intensity in the window. Intensities were extracted using a window of +-15 TOF spectra around the ion mobility apex and +-7 data points across the retention time. We simulated slicing on the first two precursor isotopes (C12 and C13) by transforming the precursor peak with the quadrupole calibration function into partitions. The resulting template for the expected fragment slicing in the two-dimensional retention time–ion mobility plane was correlated with all fragment mass slices, which contained a summed intensity of at least 1000 arbitrary units. For every such potential fragment, the correlation of each peak part and the corresponding template was calculated. Furthermore, the intensity ratio of the summed intensities between the sliced parts was calculated, and the absolute deviation from the expected value was stored.
All potential fragments with a mean correlation above 0.3 and a deviation from the intensity ratio of less than 3 were accepted for the pure fragmentation spectra. Fragments were then annotated with a 20 ppm mass tolerance, and the percentage of annotated and not annotated intensities was calculated as a purity metric.

NECFLSHKDDSPDLPK - 476.22mz

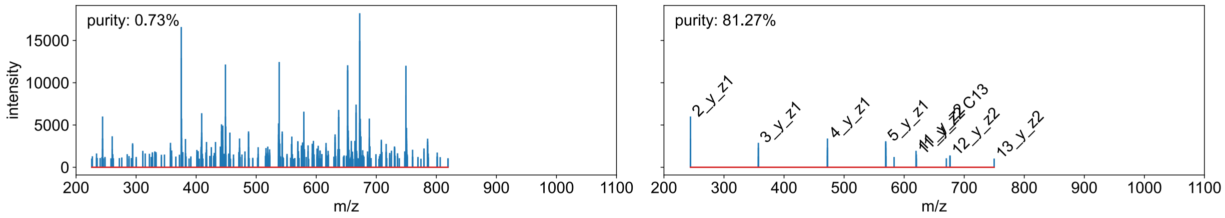


DQNYPGAIAIHHPNVAEK - 987.50mz

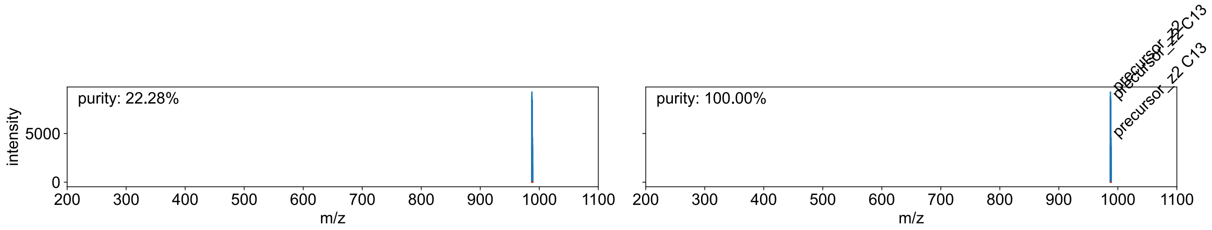


FFESFGDLSSADAILGNPK - 1007.99mz

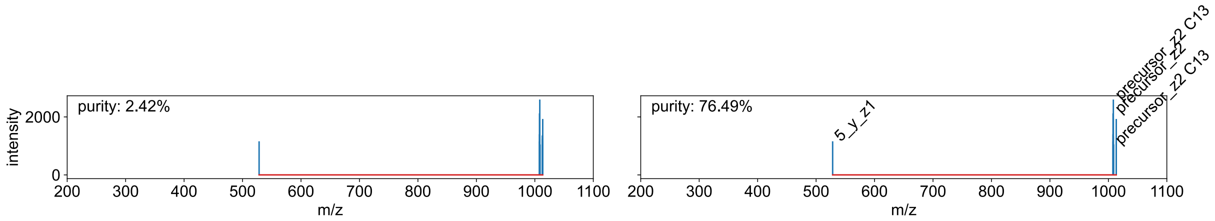


ILTATVDNANILLQIDNAR - 1034.57mz

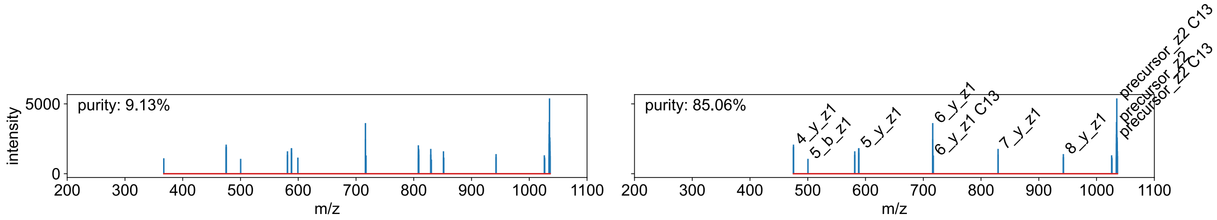


LCVLHEK - 449.74mz

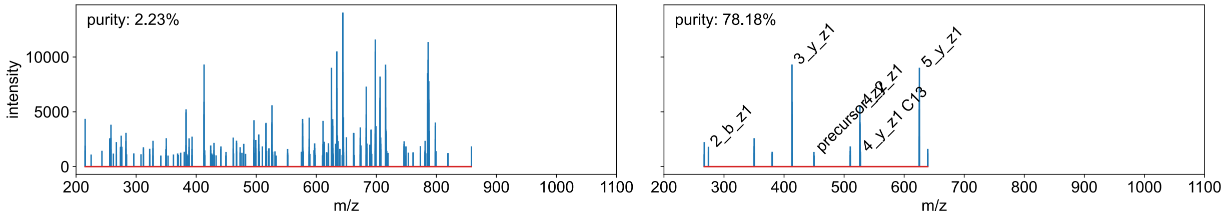


DYSQYYR - 497.72mz

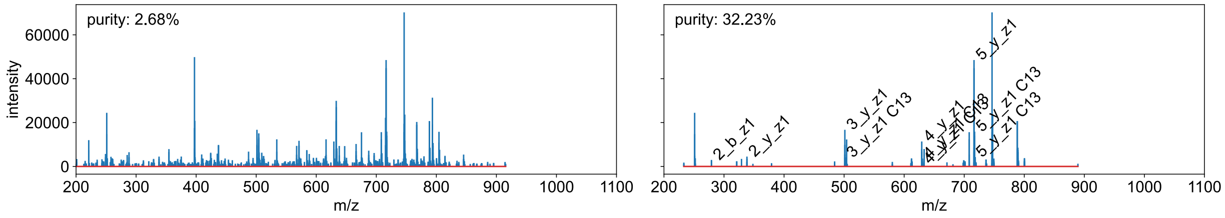

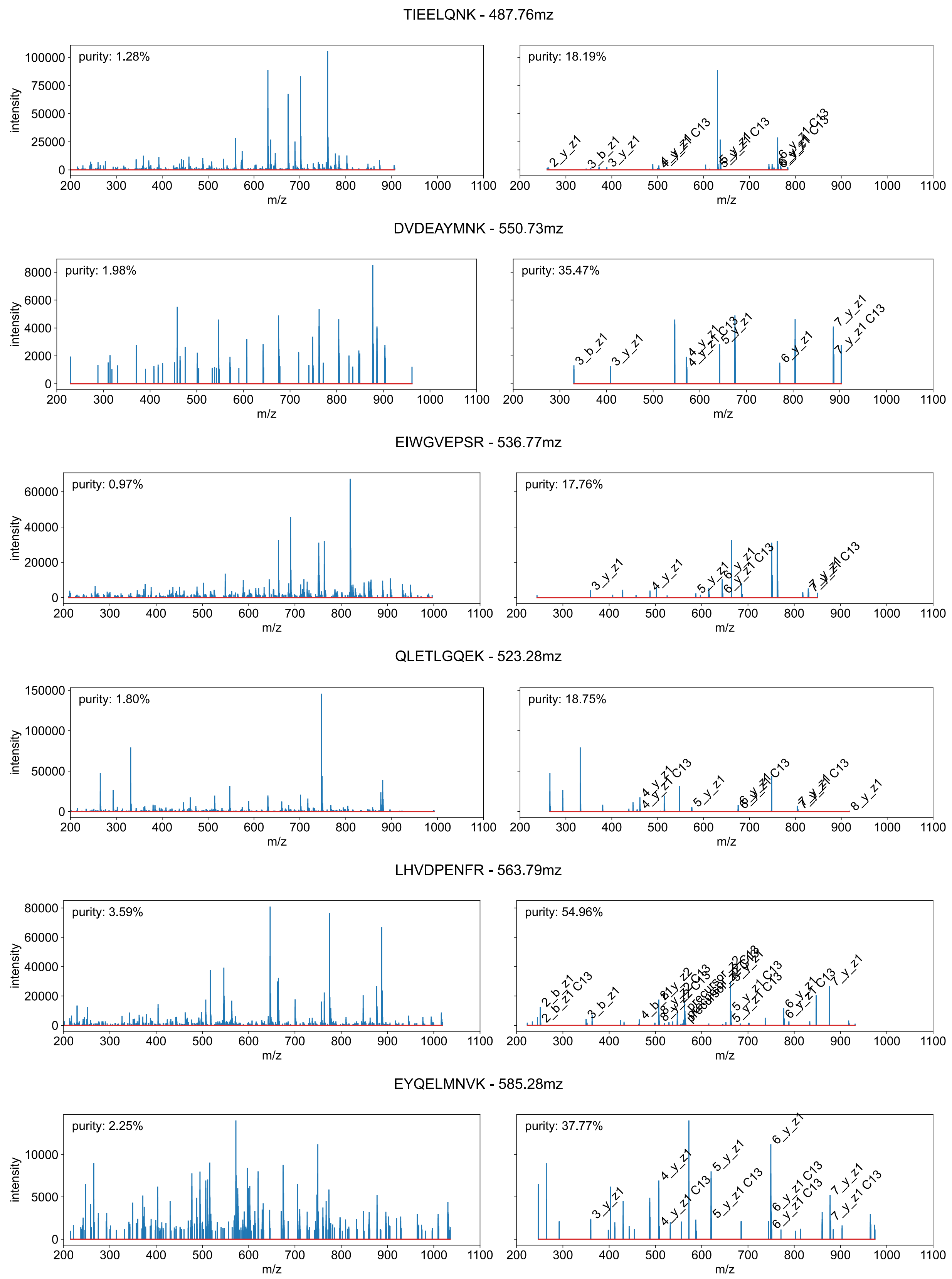
TIEELQNK - 487.76mz
purity: 1.28%
purity: 18.19%
DVDEAYMNK - 550.73mz
purity: 1.98%
purity: 35.47%
EIWGVEPSR - 536.77mz
purity: 0.97%
purity: 17.76%
QLETLGQEK - 523.28mz
purity: 1.80%
purity: 18.75%
LHVDPENFR - 563.79mz
purity: 3.59%
purity: 54.96%
EYQELMNVK - 585.28mz
purity: 2.25%
purity: 37.77%
intensity
m/z

DVDEAYMNK - 542.73mz

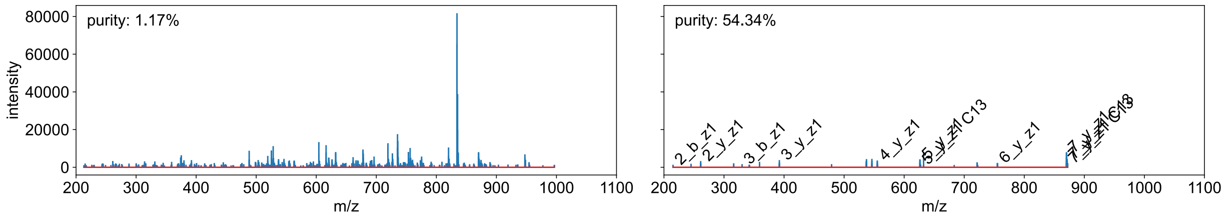


GYNAQEYYDR - 639.77mz

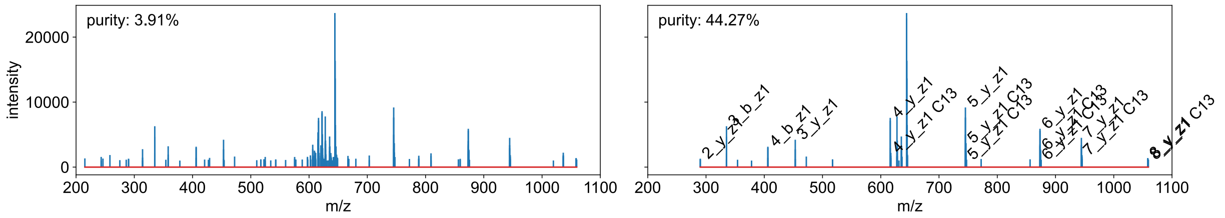


NHEEEMNALR - 621.78mz

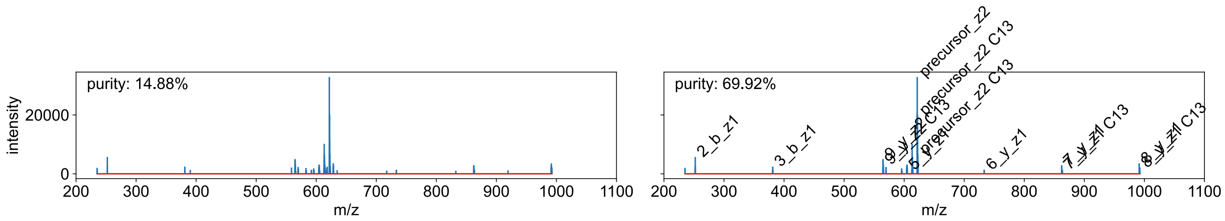


ILGEELGFVK - 552.82mz

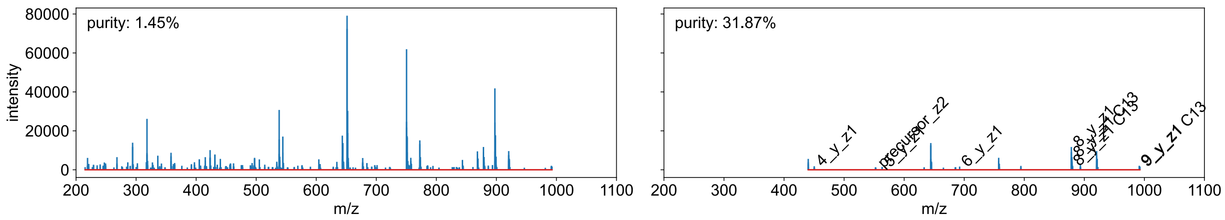


ELPDPQESIQR - 656.33mz

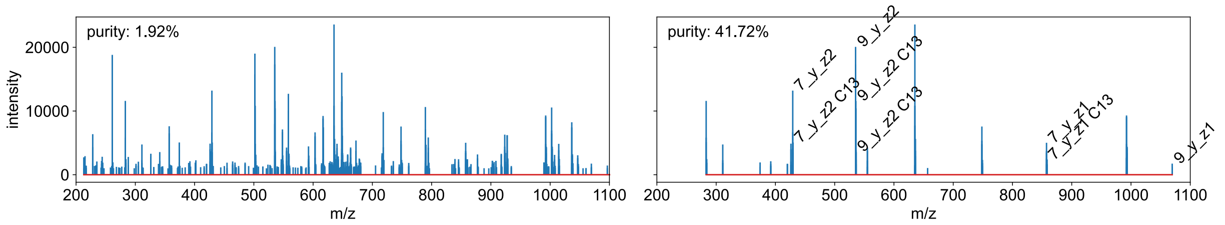


SLHTLFGDELCK - 473.90mz

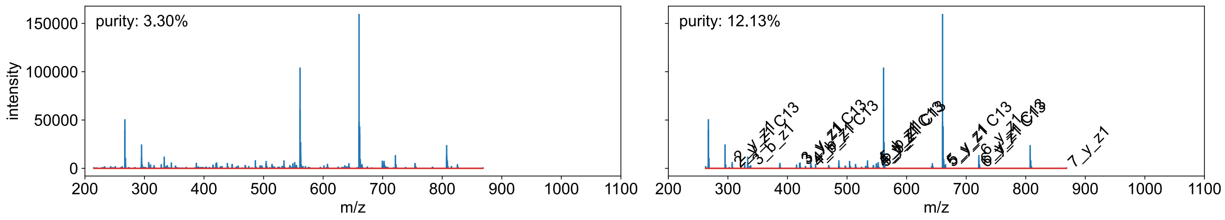

LKECCDKPLLEK - 766.89mz

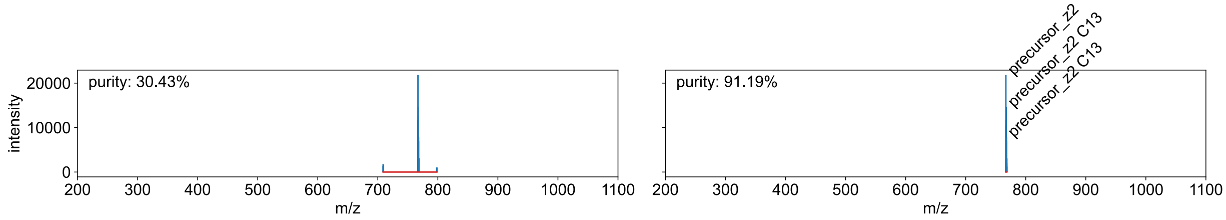


ETYGDMADCCEK - 739.77mz

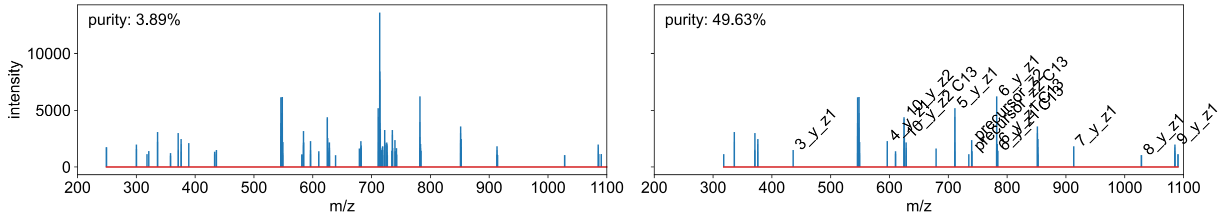


TSDANINWNNLK - 695.34mz

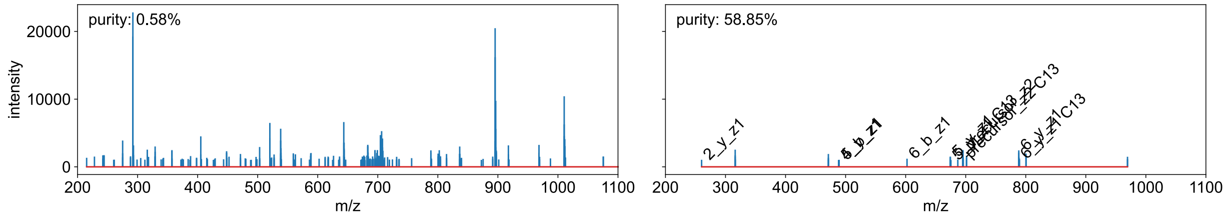


RTEMENEFVLIK - 508.93mz

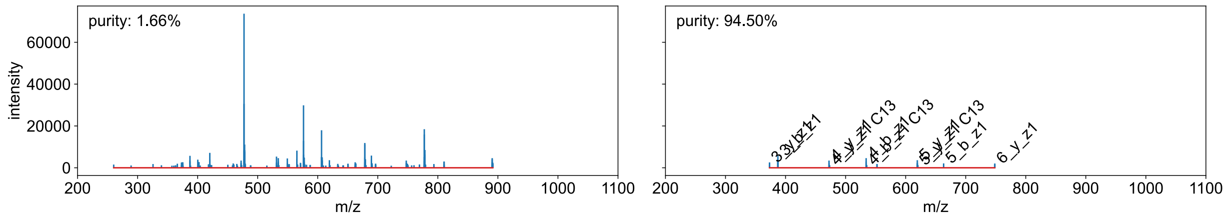


TYDSYLGDDYVR - 733.83mz

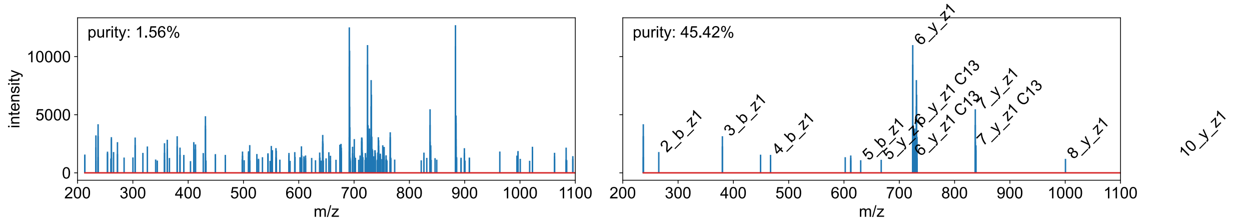


ATDAEADVASLNR - 666.82mz

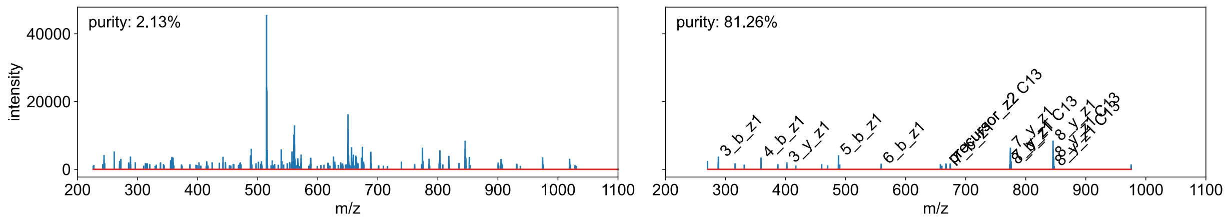

TCVADESHAGCEK - 488.53mz

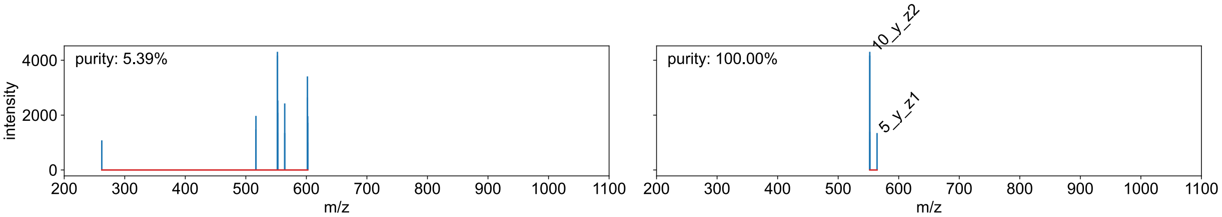


ASLEAAIADAEQR - 448.90mz

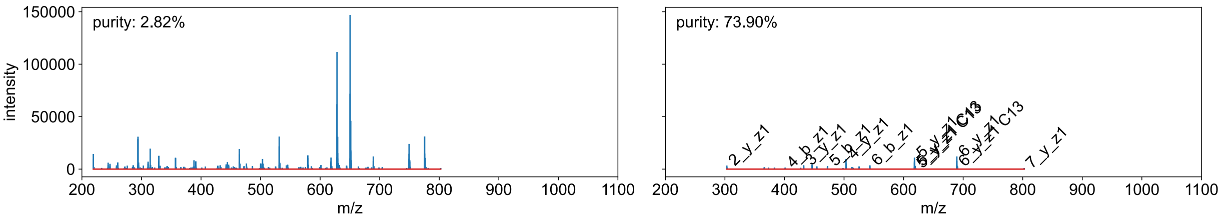


SNMDNMFESYINNLR - 940.40mz

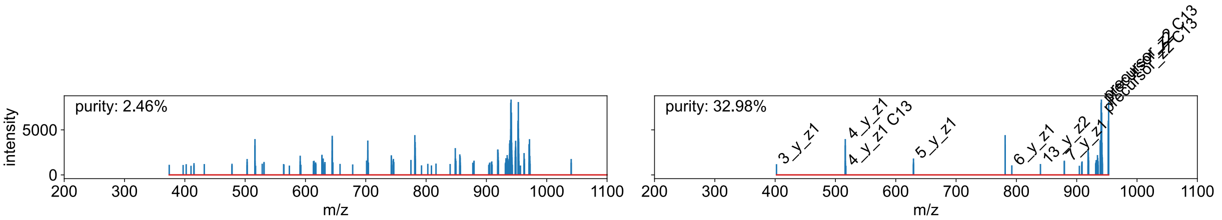


FGSEFSPELQASFQK - 851.41mz

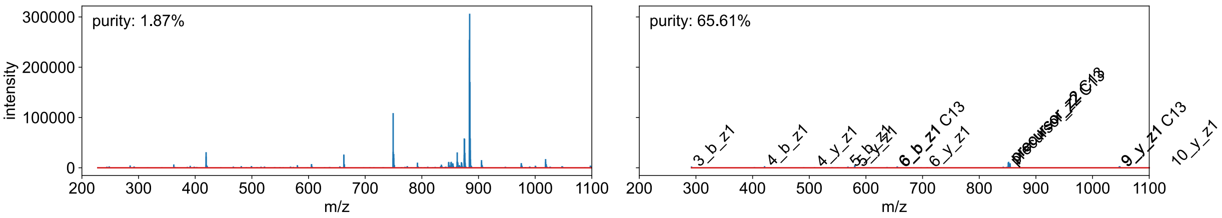


TYFPHFDLSHGSAQVK - 611.97mz

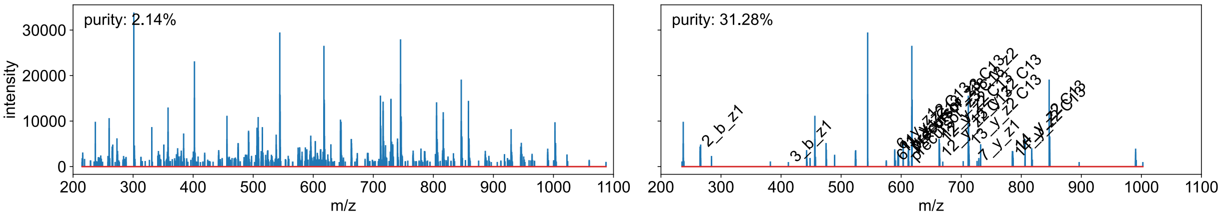
